# Bioprospecting the solar panel microbiome: high-throughput screening for antioxidant bacteria in a *Caenorhabditis elegans* model

**DOI:** 10.1101/423731

**Authors:** Kristie Tanner, Patricia Martorell, Salvador Genovés, Daniel Ramón, Lorenzo Zacarías, María Jesús Rodrigo, Juli Peretó, Manuel Porcar

## Abstract

Microbial communities that are exposed to sunlight typically share a series of adaptations to deal with the radiation they are exposed to, including efficient DNA repair systems, pigment production and protection against oxidative stress, which makes these environments good candidates for the search of novel antioxidant microorganisms. In this research project, we isolated potential antioxidant pigmented bacteria from a dry and highly-irradiated extreme environment: solar panels. High-throughput *in vivo* assays using *Caenorhabiditis elegans* as an experimental model demonstrated the high antioxidant and ultraviolet-protection properties of these bacterial isolates that proved to be rich in carotenoids. Our results suggest that solar panels harbor a microbial community that includes strains with potential applications as antioxidants.

## Introduction

Antioxidants are molecules that can protect cells against oxidative stress. For example, they can play a protective role against the biological damage derived from an excessive cellular production of reactive oxygen species (ROS). ROS are unstable metabolites of molecular oxygen (i.e. superoxide radical, hydroxyl radical or hydrogen peroxide) that are constantly generated in the cells as by-products of normal aerobic metabolism, but whose levels can increase under certain stress situations (for example, alcohol consumption, smoking or exposure to environmental pollutants) and become harmful for the cell (Al-Gubory, 2014; Rahal *et al*., 2014; Zorov *et al*., 2014; Chen *et al*., 2015). In humans, chronic oxidative stress has been associated on many occasions with the initiation and progression of a variety of diseases, including Alzheimer’s and cardiovascular diseases (such as hypertension and atherosclerosis) or cancer (Chen and Zhong, 2014; Milkovic *et al.*, 2014; Dandekar et al., 2015; Siti *et al*., 2015).

The discovery of new antioxidants from natural sources (i.e. plants or microorganisms) is of high interest for the pharmacological and food industries (Finley *et al*., 2011; Lin *et al*., 2014). The search for novel natural molecules with biotechnological applications is known as bioprospecting and, in the past, microorganisms have proved to be rich sources of natural products that have been used for the fabrication of commercial products (antibiotics, probiotics, sustainable agriculture, fermentation processes, etc.) with a wide range of applications (Mahajan *et al*., 2012; Kanchiswamy *et al*., 2015; Katz and Baltz, 2016; Choudhary *et al*., 2017; Gupta and Bajaj, 2017). Microorganisms living in harsh environments typically exhibit strategies to cope with the environmental stresses they are exposed to. In the case of microbial communities exposed to sunlight (i.e. to radiation and desiccation), these adaptations include efficient DNA repair systems, pigment production and protection from oxidative stress (Lebre *et al*., 2017), suggesting that highly-irradiated environments may be good sources of novel antioxidant-producing microorganisms. In fact, tolerances to desiccation and radiation are mechanistically correlated (Mattimore and Battista, 1996; Ragon *et al*., 2011; Slade and Radman, 2011), particularly through protection strategies against protein oxidation (Fredrickson *et al*., 2008; Fagliarone *et al*., 2017). For these reasons, in the present research we selected a highly-irradiated environment as a potential source of antioxidant-producing microorganisms: solar panels. Solar panels are man-made structures that are exposed to desiccation and high amounts of solar radiation. These harsh conditions shape the surface-inhabiting microbiome towards a highly diverse microbial community with many drought-, heat- and radiation-resistant bacteria (Dorado-Morales *et al*., 2016; Tanner *et al*., 2018). The cultivable microorganisms isolated from solar panels typically display red, orange or yellow pigmentation, which is assumed to be linked to the production of carotenoids (CRTs), natural pigments that may play a role in the protection of these microorganisms against harmful ionizing radiation and oxidative stress (Britton, 1995; Sandmann *et al*., 2015; Dorado-Morales *et al*., 2016).

Taking into account the need of screening a large number of pigment-producing bacteria isolated from the solar panels, *Caenorhabditis elegans* was chosen as an experimental organism, as it is suitable for these high-throughput screenings. *C. elegans* is a nematode which has previously been used for testing potential antioxidant compounds such as selenite (Li *et al*., 2014), cocoa products (Martorell *et al*., 2013), tyrosol (Cañuelo *et al.* 2012), or CRTs such as astaxanthin (Yazaki *et al*., 2011) or β-carotene (Lashmanova *et al*., 2015). The use of *C. elegans* as an experimental model has many advantages, such as the low cost, simplicity and quickness of the methods. Nevertheless, there is one more advantage that is of particular interest in this study: the fact that this nematode is naturally a bacteria eater, worms can directly be fed with selected bacterial strains. Laboratory *C. elegans* have a basal diet of *Escherichia coli*, but it is possible to supplement the growth medium with many ingredients of interest, including other bacteria, in order to analyze their biological activity. This functional screening method has previously been used in order to identify new antioxidant probiotic strains, such as *Lactobacillus rhamnosus* CNCM I-3690 strain (Grompone *et al*., 2012) or *Bifidobacterium animalis* subsp. *lactis* CECT 8145 strain (Martorell *et al*., 2016).

The research we present here aimed at establishing a collection of pigmented bacteria isolated from solar panels in order to select those with promising biological activities as antioxidants. For this, bacterial isolates with no record of opportunistic infections were subjected to a high-throughput antioxidant screening in *C. elegans* using the high-throughput WormTracker^™^ (WT) detector in order to quantify the survival of the worms after the addition of hydrogen peroxide to the medium. Isolates with the highest antioxidant activity were then selected for further characterization through oxidative stress and UV-protection assays. Finally, a preliminary identification of the carotenoids from the selected isolates was performed. This is the first study focused on bioprospecting the solar panel microbiome aiming at obtaining microorganisms with high potential as antioxidants, and could contribute to answer a key microbial ecology question: is there a direct link between the harshness of the studied environment and the biotechnological potential of the inhabiting microbial community?

## Results

### Isolation of pigmented bacteria

Culturing of the solar panel samples on Luria-Bertani medium (LB), Reasoner’s 2A (R2A) agar and Marine Agar (MA) medium yielded a high number of colony-forming microorganism, many of them displaying red, orange or yellow pigmentations, as previously described (Dorado-Morales *et al*., 2016). A total of 87 isolates were selected, obtained in pure culture, cryo-preserved in 20 % glycerol and subjected to taxonomic identification through 16S rRNA sequencing, with 63 isolates being successfully identified and comprising a wide range of genera and species **(Table 1)**.

**Table 1.**
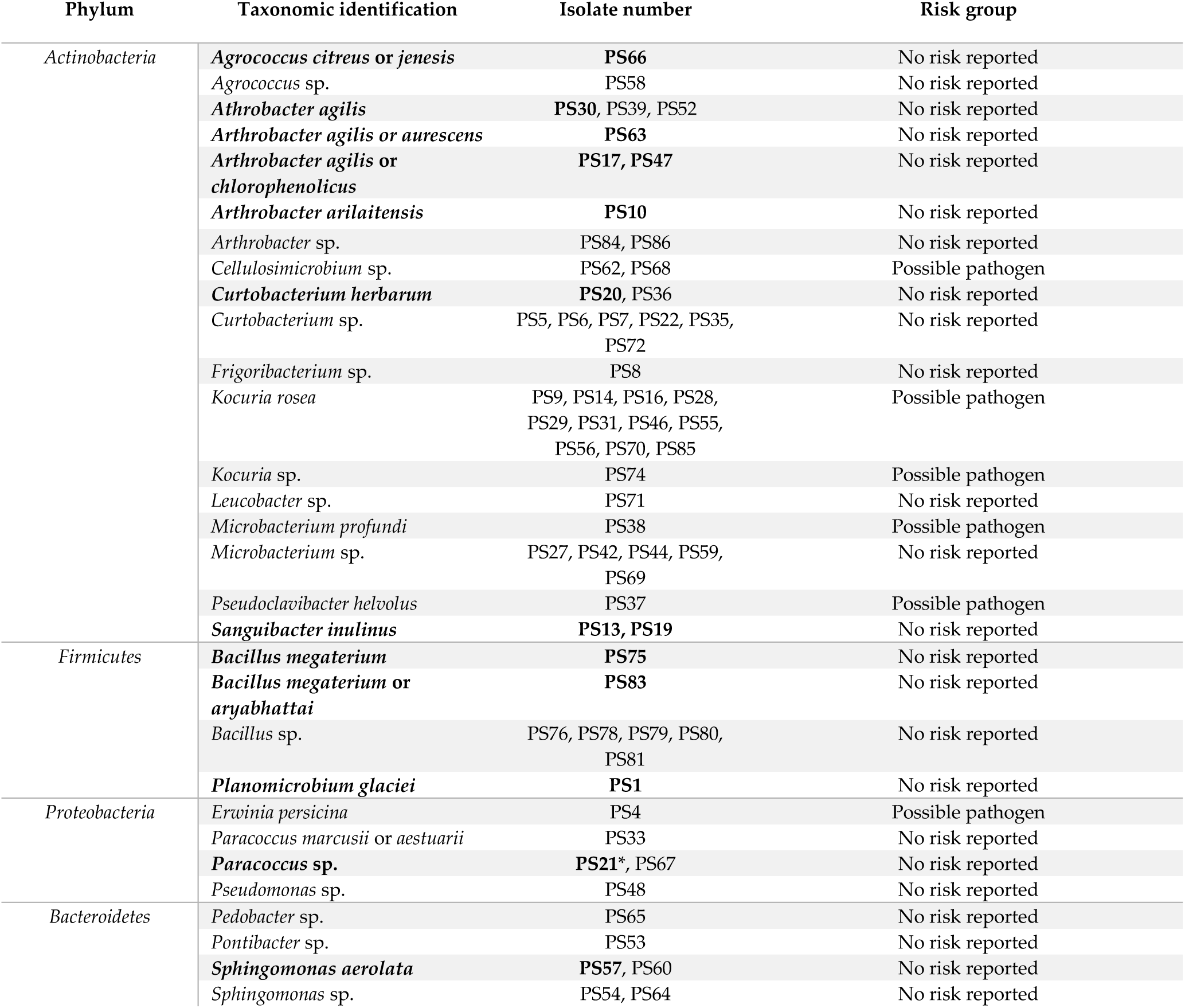
List of cultivable bacteria isolated from solar panels. Isolates fully identified at species level and with no previous reports of opportunistic infections (or species identified ambiguously with neither of the two options being opportunistic pathogens) appear in bold and were selected for further assays in *C. elegans*.

### Oxidative stress assays

After identification, only isolates fully identified at species level and with no record of opportunistic infections were selected for biological activity assays in *C. elegans*. For example, *Erwinia persicina* was not selected for these assays due to its capacity of infecting plants, causing chlorosis and necrosis in leaves (González *et al*., 2007). Isolates from the *Kocuria* genus were not selected due to increasing incidence of different types of *Kocuria* infection, mostly in immunocompromised hosts or hosts with severe underlying diseases, causing infections such as peritonitis, bacteremia or endocarditis (Purty *et al*., 2013). Finally, isolation from clinical specimens of bacteria from the genera *Microbacterium*, *Cellulosimicrobium* and *Curtobacterium* have been reported, therefore isolates from these species were not selected for biological activity assays (Francis *et al*., 2011; Gneiding *et al*., 2008; Zamora *et al*. 2018).

The selected isolates for high-throughput biological assays were the following: *Planomicrobium glaciei* (PS1), *Bacillus megaterium* or *aryabhattai* (PS83), *B. megaterium* (PS75), *Paracoccus* sp. (PS21), *Curtobacterium herbarum* (PS20), *Sanguibacter inulinus* (PS13 and PS19), *Arthrobacter agilis* (PS30), *Arthrobacter agilis* or *chlorophenolicus* (PS17 and PS47), *Arthrobacter agilis* or *aurescens* (PS63), *Arthrobacter arilaitensis* (PS10), *Agrococcus citreus* or *jenensis* (PS66) and *S. aerolata* (PS57). All these isolates were individually tested with an oxidative stress assay using the WT device, which is able to automatically assess survival of the worms through the detection of omega bends and reversals in the worm’s locomotion (Huang *et al*., 2006). Survival under oxidative stress conditions was measured after incubation of the worms for three days on Nematode Growth Medium (NGM) supplemented with each bacterial isolate at OD_600_ of 30 or of 60. Isolate PS57 did not grow well in liquid culture and was therefore discarded from the assay.

After three days of incubation with the selected pigmented isolates, some worms had not reached young adult phase and were de-synchronized. Specifically, this was the case of worms incubated with PS66, PS47, PS19 and PS20. The most extreme case was PS66, so this one was not measured in the WT device. Nevertheless, PS47, PS19 and PS20 displayed only slight differences in growth and were therefore tested. Worm activity after oxidative stress was best measured at 30 minutes after addition of hydrogen peroxide to the medium, as it is at this point when larger differences could be observed between the positive and negative controls **(Figure 1)**.

**Figure 1.**
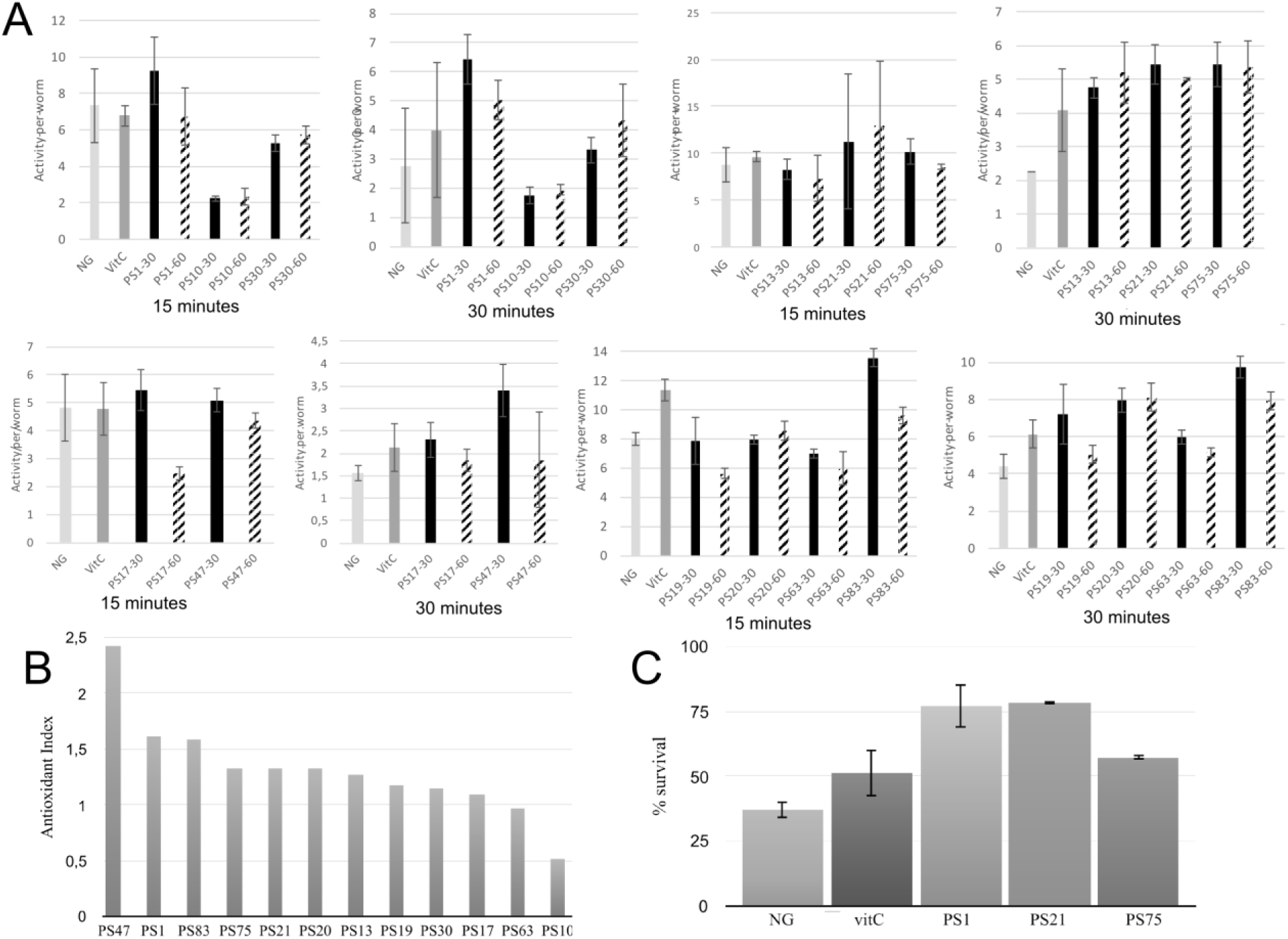
**(A)** Oxidative stress assays of the selected isolates using the WT device. Survival rate is represented in the Y-axis in the form of activity per worm, and results are shown after 15 and 30 minutes of incubation with hydrogen peroxide. **(B)** Antioxidant index (AI) of the pigmented bacterial isolates from solar panels. AI was calculated by dividing the highest activity average (at an OD600 of 30 or 60) of each isolate by the average activity of the positive control (vitC) after 30 minutes of incubation with hydrogen peroxide. **(C)** Manual oxidative stress assay results. Y-axis indicates percentage of survival of the worms after 5 hours of incubation in nematode growth medium supplemented with 20 mM hydrogen peroxide. NG (Nematode growth), negative control. VitC (vitamin C), positive control.

In general, there was no significant differences in antioxidant activity between the worms incubated with the isolates at an OD of 30 or of 60 **(Figure 1A)**, although a lower OD was beneficial for worm movement and, therefore, was the OD of choice for further experiments. After 30 minutes of incubation, PS30 did not display significant differences in activity per worm in comparison to the negative control, and PS10 displayed lower mobility than the negative control, indicating more worm mortality. On the other hand, incubation of the worms with PS1, PS13, PS21, PS75, PS17, PS47, PS19, PS20, PS63 and PS83 resulted in a higher protection of these worms against oxidative stress, with significant differences with respect to the negative control, and in some cases, with significantly higher protection in comparison to the positive control **(Figure 1A)**. In order to compare all experiments, an antioxidant index (AI) was calculated for each isolate by dividing the average activity per worm at 30 minutes when incubated with the isolate at OD 30 or 60 (the highest activity was used) by the average activity per worm of the positive control **(Figure 1B)**. Nine out of the ten tested isolates displayed higher antioxidant activity than the positive control (AI > 1), although three of these (PS47, PS19 and PS20) could not be compared to the rest due to the worms being smaller and, in some cases, not correctly synchronized.

The WT is a device that measures survival of the worms through their mobility, although this is not the most precise way to measure survival due to the fact that worms tend to have reduced mobility in liquid culture in comparison to solid medium. Therefore, the device may detect false negative results. For this reason, the best isolates according to results with the WT were selected for further, in depth characterization with the manual oxidative stress assay in order to confirm the results. Specifically, PS1, PS75 and PS21 were selected. For the manual assays, oxidative stress is applied to 5-day old adult worms instead of young adult worms, in accordance with the protocol described by Martorell *et al*. (2013).

Incubation with hydrogen peroxide resulted in a survival of approximately 37% of the worms grown on NGM with *E. coli*, whereas the survival of worms grown on NGM with *E. coli* supplemented with vitamin C (vitC) was higher, with approximately 51% survival **(Figure 1C)**, confirming the antioxidant effect of the positive control (vitC). Furthermore, the selected isolates also displayed a high antioxidant effect: incubation with PS75 resulted in around 57% survival, whereas incubation with PS1 and PS21 resulted in a survival rate of as much as 78%. These results confirm that isolates PS1, PS21 and PS75 confer a very high protection against oxidative stress in *C. elegans* and, therefore, validate the WT protocol that was designed for this project.

### UV-protection assays

The photo-protective effects of the isolated pigmented bacteria were tested *in vivo* in *C. elegans* using a UV-protection assay **(Figure 2)**.

**Figure 2.**
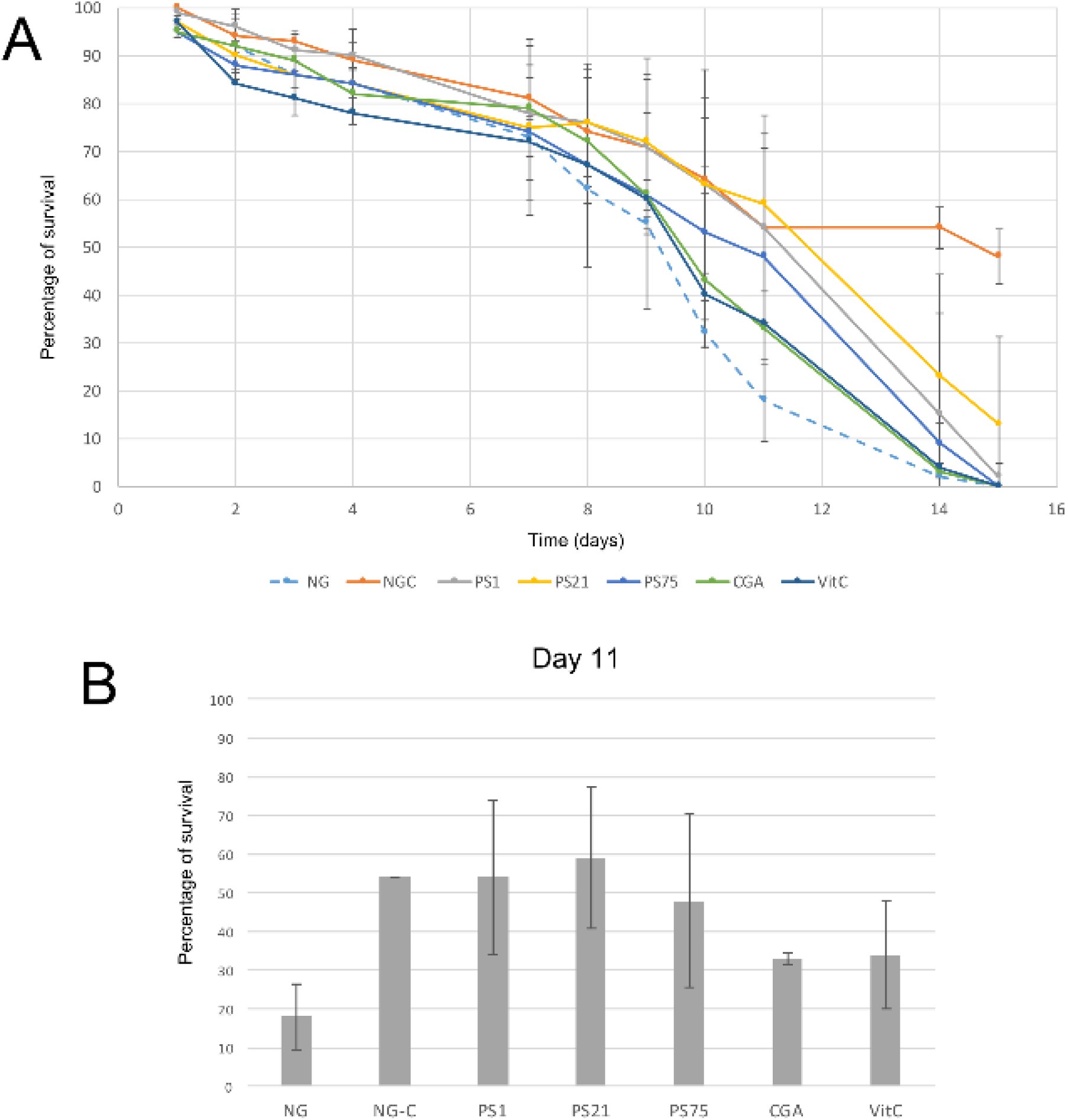
UV-light protection assay. **(A)** Y-axis indicates percentage of survival of *C. elegans* irradiated with UV-light for 45 seconds every day over a period of 15 days (X-axis). NG-C indicates the non-irradiated controls: the basal survival rate of the worms over the 15-day period. NG refers to the negative control: worms incubated in NGM with no supplements and irradiated during the 15 days. CGA and VitC are two positive controls: worms incubated with antioxidant compounds (chlorogenic acid and vitamin C) and irradiated during 15 days. Finally, PS1, PS21 and PS75 (*Planomicrobium glaciei*, *Paracoccus* sp. and *Bacillus megaterium*, respectively) indicate worms incubated with pigmented solar panel isolates and irradiated over the 15-day period in order to test the protective effect of these isolates against UV-light. **(B)** Results at day 11, in which the largest differences between the negative control and the worms fed with the pigmented isolates were observed.

There was a natural decrease in survival rate over time in the non-irradiated control (NG-C), with a survival rate at day 14 of 54% **(Figure 2A)**. Despite a general decrease of survival rate over the first 9 days **(Figure 2A)**, day 11 showed the largest decrease of the negative control survival rate (worms grown on NGM with *E. coli* and subjected to irradiation) in comparison to the survival rate of the positive controls and of the worms fed with the selected isolates **(Figure 2B)**. Worms fed with PS1 and PS21 displayed a survival rate of around 55 % at day 11, suggesting that these isolates are able to confer resistance against UV irradiation. On the other hand, although PS75 is also able to confer protection to UV-light, the survival rates are lower than the ones obtained with PS1 and PS21 (**Figure 2A)**. These results correlate with the previous ones regarding effectiveness of the strains in protecting *C. elegans* against oxidative stress: PS1 and PS21 are the isolates which confer the highest resistance, followed by PS75.

### Preliminary characterization of the carotenoid content of selected isolates

The three selected isolates (PS1, PS21 and PS75) were further studied in two different types of bacterial culture: liquid culture and solid culture. For this, pigments were extracted and analyzed by HPLC-PDA. The resulting chromatogram of each sample, together with examples of characteristic absorption spectra for CRTs peaks can be seen in **Supplementary Figure 1**. For each sample, the peaks with a characteristic CRT spectrum were integrated at their maximum wavelength and, if possible, their probable identities were assigned according to the absorbance spectrum and retention time compared to commercial standards or reported in similar chromatographic conditions. Peaks with a characteristic CRT spectrum but without assigned identity were reported as ‘not identified’ (NI) in the profile description. Peaks were quantified by interpolating the area of the peaks into calibration curves, as explained in material and methods. The relative abundance of each carotenoid can be seen in **Figure 3**, and all details (identification, peaks, maximum wavelengths, numeric indication of the spectral shape, and quantification) of the CRTs tentatively identified (TI) of each sample can be found in **Supplementary Table 1**.

**Figure 3.**
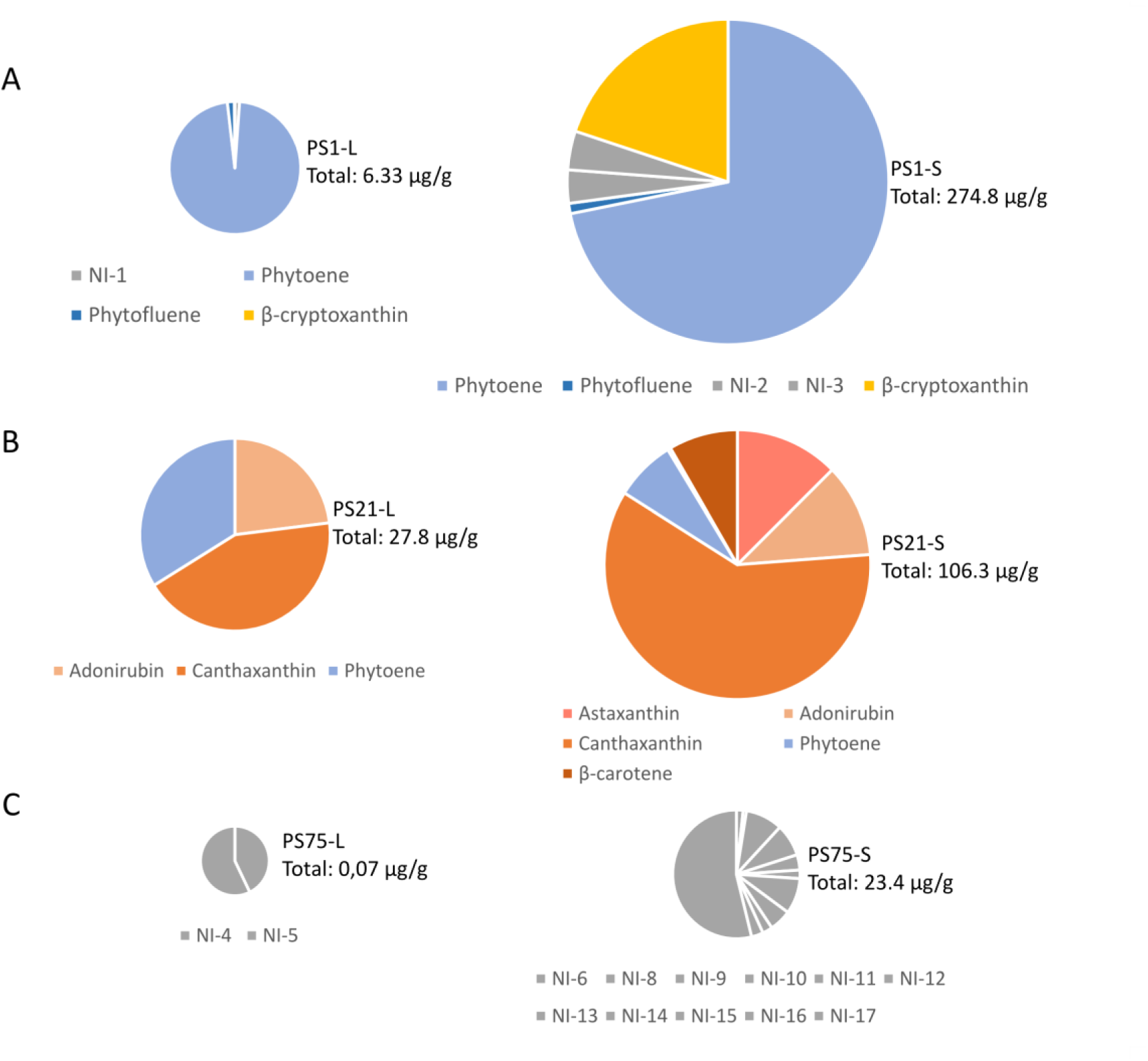
Tentative identification and quantification of the carotenoid content from the three selected isolates (A - PS1, B - PS21, C-PS75) after harvesting from liquid (L) or solid (S) culture. The total amount of CRTs is indicated next to each chart in |pg per gram of cellular pellet (dry pellet in the samples harvested from solid culture, and wet pellet in the samples harvested from liquid culture). Further details on the concentration of each carotenoid can be found in Supplementary Table 1.

## Discussion

Despite the harsh conditions, solar panels harbor a wide range of pigmented bacteria that are also shared by other harsh environments. *Microbacterium profundi* is characteristic of marine environments, isolated for the first time from deep-sea sediments of polymetallic nodule environments of the East Pacific (Wu *et al*., 2008). Other isolates are characteristic of polar environments, such as *P. glaciei*, a psychrotolerant bacterium that was first isolated from a glacier in China (Zhang *et al*., 2009), *A. agilis* (Brambilla *et al*. 2001) or *S. aerolata* (Busse *et al*., 2003); and others are characteristic of soil environments, such as *P. agri* (Roh *et al*., 2008) or many species of the *Frigoribacterium* (Kämpfer *et al*., 2000; Dastager *et al*., 2008), *Arthrobacter* (Park *et al*., 2014; Siddiqi *et al*., 2014) and *Curtobacterium* genera (Kim *et al*., 2008). Pigmentation of the bacterial isolates may play a protective role in their survival in environments with extreme temperature fluctuations and subjected to large amounts of irradiation. An intensification in the pigmentation was observed after the plates were incubated in the refrigerator for several days. Previous studies suggest that pigments such as CRTs not only play an important role in radiation protection but also in cryprotection (Dieser *et al*., 2010) due to their ability to modulate membrane fluidity in bacteria when grown under low temperature conditions (Jagannadham *et al*., 2000).

Oxidative-stress assays with *C. elegans* revealed the antioxidant properties of these isolates, making them of great interest for the pharmacological and food industries: extracts of these isolates or even the bacteria themselves could be used as promising treatments for conditions in which oxidative stress plays an important role. On the other hand, the UV-protection assays suggest that the pigmented bacteria isolated from solar panels could also play a protecting role in this type of stress, which is of high interest for the cosmetic industry, specifically in the fabrication of products that protect against sunlight-induced skin damage. The three isolates selected for UV-light protection assays due to the promising results obtained in the oxidative-stress tests (*P. glaciei* or PS1, *Paracoccus* sp. or PS21 and *B. megaterium* or PS75) were further tested through HPLC-PDA analysis to shed light on their CRTs composition.

*P. glaciei* (PS1) was first described by Zhang *et al*. (2009), who indicated that it displayed yellow-to-orange pigmentation. Our results suggest that the main CRTs present in PS1 may be phytoene and β-cryptoxanthin, and previous studies have demonstrated the antioxidant and free radical scavenging properties of β-cryptoxanthin, phytoene and phytofluene (Martínez *et al*., 2014; Ni *et al*., 2014). It would be interesting to consider whether the high antioxidant capacity of this isolate could be related to the presence of phytoene together with the colored CRT, β-cryptoxanthin.

Interestingly, although *Paracoccus* sp. (PS21) harvested from liquid culture was seen to be rich in pigments probably corresponding to adonirubin (TI), canthaxanthin and phytoene, when harvested from solid medium CRT composition included also astaxanthin (T) and β-carotene. The CRTs present in PS1 and PS21 could be commercially valuable as they have many applications (Sandmann, 2015): β-carotene and canthaxanthin are used as food colorants and feed additives, especially in aquaculture, whereas astaxanthin and phytoene are widely used in the cosmetic industry.

Finally, PS75 was identified as *B. megaterium*, a spore-forming species (Mitchell *et al*., 1986). Although no identity was assigned to CRT peaks in PS75 extracts, the absorbance spectrum and retention time in the used chromatographic conditions of NI-17 and other minor peaks (NI-14 to -16) in solid culture, and NI-4 and 5 in liquid culture, are compatible with methyl esters of glycosyl-apo-8’-lycopene, orange colored derivatives of a C30 apo-8’-carotenoid pathway that occurs in certain *Bacillus* species (Pérez-Fons *et al*., 2011). Moreover, the NI-6 to NI-13 compounds and phytoene-like may also correspond to glycosyl-3-4-dehydro-8’-apolycopene esters and apo-8-phytoene which have been identified vegetative cells and spores of *Bacillus* spp. species (Pérez-Fons *et al*., 2011). In relation to the oxidative stress and UV-resistant assays, this isolate had less antioxidant activity in comparison to PS1 and PS21.

In conclusion, after selecting a number of pigmented isolates from solar panels according to their low biological risk and testing them *in vivo* in order to elucidate their biological activity, nine out of the ten selected isolates displayed a higher antioxidant activity than the positive control. The isolates with highest antioxidant activity, PS1 (*P. glaciei*), PS21 (*Paracoccus* sp.) and PS75 (*B. megaterium*) were validated with a manual oxidative stress assay, confirming the previous results and validating the protocol designed and used for oxidative stress assay in the WT device. Furthermore, the three selected strains also displayed UV-protection properties, with values once again higher than the positive control in the case of PS1 and PS21. The high antioxidant properties of these isolates are promising from a pharmacological point of view. Specifically, extracts of these bacteria or artificial combinations of their active compounds, could be useful for the design of new treatments against diseases in which oxidative stress plays a crucial role.

Taken together, our results provide new data on the biological activity of bacterial strains from solar panels with very high antioxidant and UV-protection properties. This is the first report describing the biotechnological potential of pigmented bacterial strains from solar panels using a *C. elegans*-based model.

## Methods

### Sampling

Samples were collected from six solar panels located on the rooftop of the Faculty of Economics of the University of Valencia on the 30th November 2015. Sampling was performed by washing the solar panels with sterile Phosphate-Buffered Saline (PBS) and by scraping the surface with sterile glass wipers as previously described (Dorado-Morales *et al*., 2016). The resulting liquid was collected using sterile pipettes and stored in 50 mL Falcon tubes, which were then transported to the laboratory on ice, where cultivation, isolation and identification of the strains was performed.

### Cultivation and isolation of pigmented bacterial strains

Solar panel samples were cultivated on LB agar, R2A agar (Reasoner *et al*., 1985) and MA by spreading 50 μL of the collected liquid to each plate. Then, samples were left to settle for 30 minutes, allowing the larger sized particles - including many fungi - to sediment, and 50 μL of the supernatant were plated on LB, R2A agar and MA. By allowing the samples to settle, fungal growth was reduced when cultivating the samples on the different culture media. Plates were incubated at room temperature for one week.

After one week of incubation, colonies were selected based on colony morphology (colour, size, texture, etc.) and isolated in pure culture by re-streaking on fresh medium. These pure isolates were conserved at -80^°^C in 20 % glycerol for future use.

### 16S rRNA sequencing

For 16S rRNA sequencing, a 500-bp fragment of the hypervariable region V1-V3 of the isolates was amplified by colony PCR, using universal primers 28F (5’-GAG TTT GAT CNT GGC TCA G-3’) and 519R (5’-GTN TTA CNG CGG CKG CTG-3’). Isolates whose 16S rRNA failed to amplify from colony templates were amplified again with the same PCR program plus an initial step of incubation for 10 minutes at 100 ^<u>o</u>^C. Amplicons were checked in 1.4 % agarose gel and then precipitated overnight in isopropanol 1:1 (vol:vol) and potassium acetate 3M pH 5 1:10 (vol:vol). Precipitated DNA was washed with 70 % ethanol, resuspended in Milli-Q water (Merck Millipore Ltd, Tullagreen, Cork, Ireland) and quantified with a Nanodrop-1000 Spectrophotometer (Thermo Scientific, Wilmington, DE, USA). Amplicons were tagged using BigDye^®^ Terminator v3.1 Cycle Sequencing Kit (Applied Biosystems, Carlsbad, CA, USA) and sequenced with the Sanger method by the Sequencing Service (SCSIE) of the University of Valencia (Spain). The resulting sequences were manually edited using Pregap4 (Staden Package, 2002) to eliminate low-quality base calls. The final sequence for each isolate was compared to nucleotide databases using the NCBI BLAST tool.

### Oxidative stress assays with WormTracker

Experiments were carried out with the wild-type *C. elegans* strain N2 (Bristol), which was routinely propagated at 20 ^<u>o</u>^C on Nematode Growth Medium (NGM) plates supplemented with *E. coli* strain OP50 as the regular food source.

Worms were synchronized by isolating eggs from gravid adults at 20 ^<u>o</u>^C. Synchronization was performed on NGM plates with *E. coli* OP50 as a negative control, *E. coli* OP50 plus vitamin C (vitC) at 20 μg/mL as a positive control **(Supplementary Figure 2A)**, or *E. coli* OP50 plus the pigmented isolates in order to test antioxidant properties of the bacteria. The isolates were grown overnight in liquid LB medium at 28 ^<u>o</u>^C and 180 rpm, optical density at 600 nm (OD_600_) was adjusted to 30 and to 60, and 50 μL of the bacterial suspension was added to the plates. The synchronized worms were incubated for a total of three days on the previously described plates, until reaching young adult stage.

Young adult worms were collected and washed three times with M9 buffer, and finally resuspended in 100-200 μL of the buffer. Worms were then transferred by pipetting to 96-well plates (10-30 worms per well) containing M9 buffer. After transferring all the worms, hydrogen peroxide was added to the wells, reaching a final concentration of 1.2 mM of hydrogen peroxide **(Supplementary Figure 2B)**. Mobility of the worms was measured with the WT during 60 minutes (four measurements of 15 minutes), and data was collected in the form of “worm activity” (mobility of the worms). Finally, the number of worms in each well was manually counted in order to normalize the mobility data by the number of worms. All assays were performed with two biological replicates.

### Manual oxidative stress assays

Manual assays were also carried out with the wild-type *C. elegans* strain N2 (Bristol), routinely propagated and synchronized as previously described (on NGM with *E. coli* OP50 as a negative control, and supplemented with pigmented isolates at an OD_600_ of 30 for biological assays), except for the positive control, which in this case was vitC at 10 μg/mL. Young adult worms were transferred to fresh plates once every two days, until reaching 5-day adult stage. Then, these worms were transferred to plates containing basal medium supplemented with 2 mM hydrogen peroxide and incubated for 5 hours at 20 ^<u>o</u>^C. After incubation, the survival rate of the worms for each condition (negative control, positive control and fed with pigmented bacteria) was calculated by manually assessing survival of the worms. Two biological replicates were performed for every condition.

### UV-protection assays

Wild-type *C. elegans* strain N2 (Bristol) worms were synchronized on NGM plates with *E. coli* OP50 as a negative control, *E. coli* OP50 plus vitC (0.1 μg/mL) or plus chlorogenic acid (CGA) (0.1 μg/mL) as positive controls, or *E. coli* OP50 plus the pigmented isolates (50 μL of an overnight culture adjusted to OD 30) in order to test the UV light protection properties of the bacteria.

Synchronized worms were propagated for 15 days on the different types of medium, irradiated daily for 45 seconds in the laminar flow hood with UV light and transferred to new medium every two days, as previously described (Iriondo-DeHond *et al*., 2016). Survival rate of the worms was manually recorded every day and the assay was performed with biological duplicates.

### Pigment extraction

Carotenoid extraction was performed with two types of bacterial cultures: grown on solid (S) and in liquid (L) medium for one week and 12 hours (overnight), respectively. For CRTs extraction from isolates grown on solid medium, bacterial cells were collected from solid LB medium after one week of incubation at room temperature. Cells were resuspended in PBS and concentrated through centrifugation at 13000 rpm for 3 minutes. The supernatant was discarded and pellets were dried completely with a vacuum-connected centrifuge (DNA Speed Vac, DNA120, Savant). Then, dry weight was determined. For the exponential phase samples, overnight cultures of selected isolates were collected and the wet weight was determined for each sample.

Bacterial pellets were resuspended and washed in Tris-Buffered Saline (TBS) solution, and centrifuged. Pelleted cells were frozen in liquid nitrogen (N2) three times, followed by addition of methanol (Sharlau, HPLC grade) (ten times the volume of the pellet) and sonication in a XUBA3 ultrasonic water bath (35 W; Grant Instruments, Cambridge, England) for five minutes, in order to break the bacterial cells. Samples were vigorously shaken and centrifuged, and then the upper layer of colored methanol was transferred to a clean tube. This step was performed several times until a non-colored pellet was obtained.

Dicloromethane (HPLC grade) and water (Milli Q grade) (both at ten times the volume of the original pellet) were added to the methanol extract in order to separate organic and aqueous phases. Samples were vigorously shaken, centrifuged, and the aqueous phase was discarded. This step was performed twice, finally yielding CRT extracts in dicloromethane. Samples were then dried under N2 and kept at -20^°^C until analysis by HPLC-PDA. All steps were performed under dim light to avoid CRTs modifications such as photodegradation, isomerizations or structural changes.

### HPLC-PDA analysis

CRT composition of each sample was analysed by using an HPLC with a Waters liquid chromatography system (Waters, Barcelona, Spain) equipped with a 600E pump and a 2998 photodiode array detector (PDA). Empower software (Waters, Barcelona, Spain) was used for HPLC program set up and chromatogram analysis. A C_30_ CRT column (250 mm × 4.6 mm, 5 μm) coupled to a C_30_ guard column (20 mm × 4.0 mm, 5 μm) (YMC GmbH, Germany) was used. Samples were prepared for HPLC analysis by dissolving the CRT extracts in CHCl_3_:MeOH:acetone (3:2:1, v:v:v), followed by centrifugation for 2 minutes at 13000 rpm in order to discard any solid residues. CRT separation was performed with a ternary gradient elution, with an initial solvent composition of 90% methanol (MeOH), 5 % water and 5 % methyl tert-butyl ether (MTBE). Solvent composition changed during the analysis as described by Carmona *et al*. (2012) and Alquezar *et al*. (2008). After each analysis, the initial conditions were re-established and equilibrated before the next injection. The flow rate was 1 mL min^-1^ and column temperature was 25 ^<u>o</u>^C. A volume of 20 μL of each sample was injected and the photodiode array detector was set to scan from 250 to 540 nm. A Maxplot chromatogram was obtained for each sample that plots each CRT peak at its corresponding maximum absorbance wavelength.

CRTs were identified by comparison of the absorption spectra and retention times with the available standards or with data obtained in similar experimental conditions and described in the literature (Britton et al., 1998). For quantification, the chromatographic peaks of each CRT were integrated in their maximum wavelength and the resulting area of the peak was interpolated in different calibration curves that were already set up in the laboratory. The available calibration curves were: canthaxanthin (Sigma), lutein (Sigma), β-carotene (Sigma), β-cryptoxanthin (Extrasynthese). Standards of phytoene and phytofluene were obtained from peel extracts of orange fruits (Rodrigo *et al*. 2003) and HPLC purified. Quantification of adonirubin, astaxanthin and echineone was performed using the calibration curve of β-carotene, with values expressed as equivalents of β-carotene. As for the non-identified CRTs, they were quantified using either the β-carotene or the lutein calibration curves depending on their retention times and spectra.

## Acknowledgements

Financial support from the Spanish Government (grant Helios, reference: BIO2015-66960-C3-1-R co-financed by FEDER funds and Ministerio de Economía y Competitividad) and from the Regional Government of Valencia (grant MICROBIOSOL, reference: IFIDUA/2015/10 financed by IVACE) are acknowledged.

## Contributions

The project was designed by MP, JP and DR. Sampling and isolation/identification of the strains was performed by MP, JP and KT. Experiments involving *C. elegans* were performed by SG, PM, DR and KT. Characterization of the pigment content was performed by MJR, LZ and KT. Manuscript has been written and revised by all authors.

